# Understanding emergence of antimycobacterial dose dependent resistance

**DOI:** 10.1101/2022.09.02.506358

**Authors:** Frank Kloprogge, Julio Ortiz Canseco, Lynette Phee, Zahra Sadouki, Karin Kipper, Adam A. Witney, Neil Stoker, Timothy D. McHugh

## Abstract

Concentration dependency of phenotypic and genotypic isoniazid-rifampicin resistance emergence was investigated to obtain a mechanistic understanding on how anti-mycobacterial drugs facilitate the emergence of bacterial populations that survive throughout treatment. Using static kill curve experiments, observing two evolution cycles, it was demonstrated that rifampicin resistance was the result of non-specific mechanisms and not associated with accumulation of drug resistance encoding SNPs. Whereas, part of isoniazid resistance could be accounted for by accumulation of specific SNPs, which was concentration dependent. Using a Hollow Fibre Infection Model it was demonstrated that emergence of genotypic resistance only occurs when antibiotic levels fall below MIC although MICs are typically maintained following clinical dosing provided that adherence to the regimen is good. This study showed that disentangling and quantifying concentration dependent emergence of resistance provides improved rational for drug and dose selection although further work on understanding underlying mechanisms is needed to improve the drug development pipeline.

**One Sentence Summary:** Disentangling and quantifying concentration dependent emergence of resistance will contribute to better informed drug and dose selection for anti-mycobacterial combination therapy.

## INTRODUCTION

Tuberculosis (TB) is caused by *Mycobacterium tuberculosis* and remains the most infectious disease caused by a single bacterium with highest mortality worldwide at 5.8 million new infections in 2020 and 1.3 million HIV-negative and 214,000 HIV-positive deaths (*1*). Treatment outcome is generally favorable for drug sensitive infections (DS-TB) as long as adherence to treatment is good. However, the treatment of TB is protracted, currently at least 6 months, and it can cause serious adverse events resulting in the need to change the treatment regimen (*2*). The development of resistant forms of TB forms a major threat to global health. In 2020, 157,842 and 25,630 laboratory-confirmed Multi Drug Resistant/Rifampicin Resistant (MDR/RR)-TB and Extensively drug-resistant (XDR)-cases were reported while treatment outcomes are generally poor for these forms of disease at 59% and 52% MDR/RR-TB and XDR-TB compared to 86% for DS-TB (*1*).

Recently, various consortia have been trying to improve TB treatment efficacy and shorten duration by development of novel anti-mycobacterial treatment combinations for DR-TB, MDR-TB or pan-TB (*3*) (NCT03338621, NCT02342886, NCT03474198 and NCT03086486). Other studies have tried to optimize standard dosing regimens by increasing the rifampicin dosage for treatment of active TB (*4*) or by increasing the isoniazid dosage for treatment of MDR-TB in children (*5*).

Despite many of these efforts showing promising interim results or proven success, treatment remains long at a minimum of four months. Meanwhile, development of further treatment shortening is hampered by the lack of a rationale for selection of candidate drug combinations entering clinical phases of research and inefficient protocols for these clinical trials. Obtaining a mechanistic understanding of how anti-mycobacterial drugs facilitate the emergence of bacterial populations that survive throughout the treatment would contribute to addressing the challenges around selecting candidate drug combinations for evaluation in clinical trials.

The causative mechanisms enabling survival of antibiotic exposure are many and include inheritable genotypic resistance, caused by Single Nucleotide Polymorphisms (SNPs) that are known to be associated with drug specific resistance and result in an increase in Minimum Inhibitory Concentration (MIC). Phenotypic resistance on the other hand is used to describe all other resistance with no known drug specific SNPs described, often referred to as antibiotic persistence and tolerance (*6, 7*) and may include non-specific mechanisms such as drug efflux or cell wall permeability.

Antibiotic tolerance refers to an increased ability of the bacterial population to survive antibiotic exposure without increase in MIC. These manifests itself in greater minimum duration to kill 99% of the population (MDK_99%_). Antibiotic tolerance is caused by a combination of general and drug specific mechanisms that are independent of antibiotic class and include reduced growth rates, metabolic shifts and increased activity of efflux pumps (*6, 7*).

Antibiotic persistence refers to a predestined sub-population of bacteria, often 0.01-1 % of an inherently heterogenous population of bacteria, that is able to survive throughout antibiotic treatment. The persistent sub-population is more difficult to kill, i.e. has a longer MDK_99%_, while the majority of the bacterial population is fully susceptible, i.e. has a shorter MDK_99%_, and this results in a characteristic bi-phasic killing profile over the course of antibiotic exposure without increase in MIC. Antibiotic persistence is not inheritable, meaning a heterogenous population will regrow when antibiotic pressure is taken off (*6-8*).

Bacteria within the second phase of a bi-phasic killing profile may consist of different sub-populations. Little research has been done in the domain of disentangling and quantifying genotypic from phenotypic resistance in the tail of bi-phasic killing profiles for antimicrobial combination therapy, even though it forms the cornerstone of anti-TB therapy.

Isoniazid and rifampicin form the backbone of standard drug-sensitive anti-tuberculosis therapy and isoniazid enhances bi-phasic bacillary killing from sputum, but concomitant therapy with rifampicin does not prevent this from happening (*8, 9*). Given the mutation rate, in H37Rv, of isoniazid (2.56 × 10^−8^ - 3.2 × 10^−7^ (*10, 11*)) and rifampicin to a lesser extent (6 × 10^−10^ - 2.4 × 10^−7^ (*11-15*)), this could be emergence of heterogenous genotypic resistance, phenotypic resistance or a combination of both. Disentangling and quantifying emergence of genotypic and phenotypic resistance over the course of isoniazid and rifampicin combination therapy can therefore inform ongoing *in-vivo* investigations evaluating the increase in rifampicin and isoniazid dose in order to achieve improved and sustained overall anti-mycobacterial activity (*4*).

To that end the concentration dependency of emergence of phenotypic and genotypic isoniazid-rifampicin resistance was investigated and the impact of *in-vivo* mimicking pharmacokinetic profiles at a range of dose scenarios were explored.

## RESULTS

### Pharmacodynamic interactions and emergence of resistance

Static kill curve experiments, in which broth cultures are inoculated and followed for two evolution cycles, were used to study emergence of bacterial sub-populations that were able to survive isoniazid and/or rifampicin exposure. Isoniazid and rifampicin drug effects disappear over the course of the experiment, whether given alone or in combination, both in the first and in the second evolution cycle.

However, when isoniazid was used alone, potency decreased (associated with an increase in IC_50_, i.e. the concentration at which half-maximum antimycobacterial effect is achieved, of 402%) after repeated exposure but a resistant population, i.e. described by the tail in bi-phasic mycobacterial killing model, emerged later in time compared to the first exposure cycle (associated with a decrease in model parameter λ, i.e. onset of susceptibility loss as a function of time, of 16.9%) (Table 1 and Figure 1A & 1C). Likewise, when studied alone rifampicin potency decreased (associated with an increase in IC_50_ of 182%) after repeated exposure but unlike the isoniazid treatment resistance emerged around the same time in the first and second evolution cycle (Table 1 and Figure 1B & 1D).

**Table 1.** Pharmacodynamic parameter estimates. Population, random between experiment variability (BSV), precision (%RSE) and overfitting (Shrinkage) estimates.

| Parameter | Population estimate<br>(95%CI) | %RSE | BSV(CV%) | Shrinkage<br>(SD)% |
| --- | --- | --- | --- | --- |
| knet | 0.368 (0.346, 0.391) | 3.14 | 15.5 | 5.84% |
| bmax | 86.6 (76.7, 97.7) | 1.39 | 32.6 | 2.06% |
| B0 | 134 (112, 160) | 1.86 | 47.7 | 5.43% |
| E<sub>MAX-INH</sub> | 0.734 (0.724, 0.745) | 2.35 | 2.34 | 87.10% |
| RIF interaction at re-exposure effect | -0.568 (-0.646, -0.491) | 6.96 |  |  |
| IC<sub>50-INH</sub> | 783 (721, 851) | 0.635 | 13.1 | 87.20% |
| Re-exposure effect | 4.02 (3.8, 4.24) | 2.79 |  |  |
| Hill<sub>INH</sub> | 0.756 (0.634, 0.901) | 32.1 | 15.9 | 85.70% |
| $\lambda_{INH}$ | 0.212 (0.21, 0.214) | 0.359 | | |
| Re-exposure effect | -0.169 (-0.169, -0.168) | 0.194 |  |  |
| RIF interaction effect | -0.32 (-0.321, -0.318) | 0.218 |  |  |
| E<sub>MAX-RIF</sub> | 0.43 (0.43, 0.43) | 0.047 | 44 | 45.90% |
| INH interaction at re-exposure effect | 0.421 (0.403, 0.438) | 2.12 |  |  |
| IC<sub>50-RIF</sub> | 46.8 (46.7, 46.8) | 0.0138 | 32.1 | 84.80% |
| Re-exposure effect | 1.82 (1.81, 1.83) | 0.352 |  |  |
| $\lambda_{RIF}$ | 0.153 (0.153, 0.153) | 0.0214 | | |
| INH interaction effect | 0.582 (0.58, 0.584) | 0.18 |  |  |
| Residual Variability (proportional) | 0.232 |  |  |  |

**Fig. 1.**
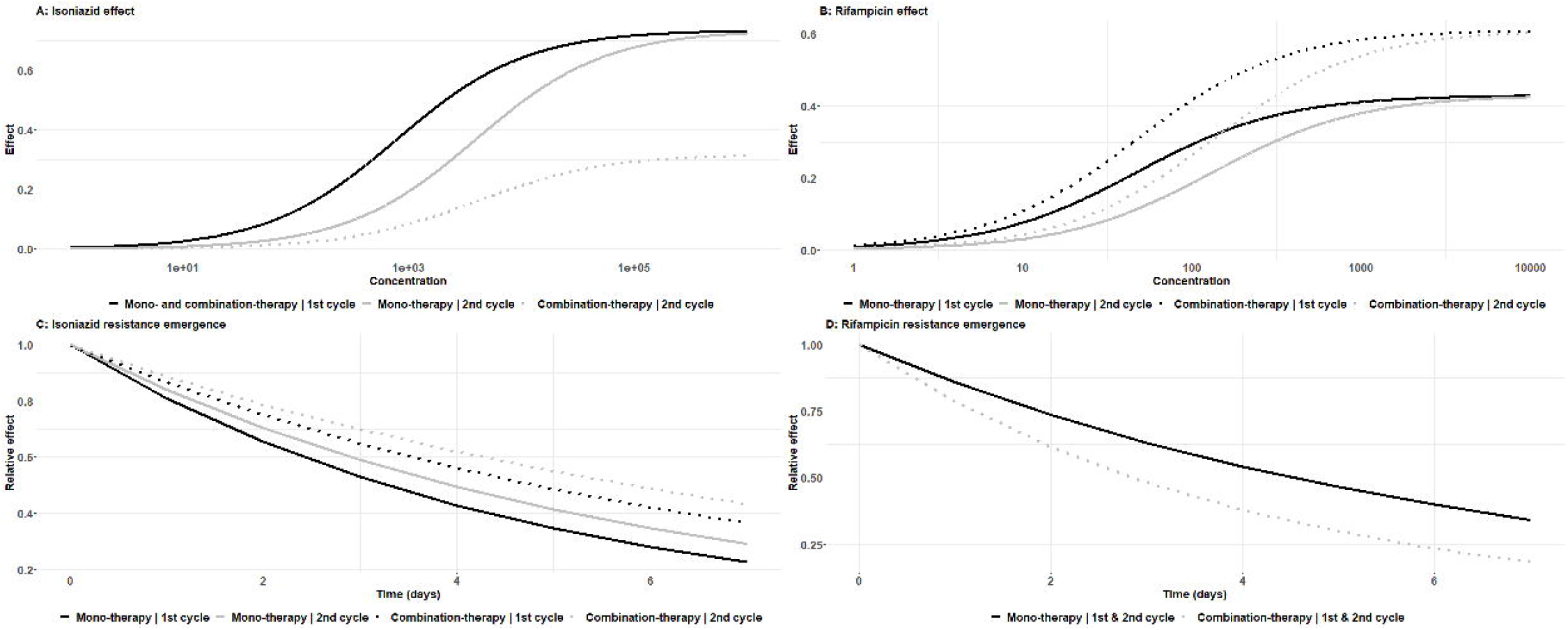
Pharmacodynamic isoniazid-rifampicin interactions. Concentration effect curves for isoniazid (**A**) and rifampicin (**B**) and emergence of phenotypic resistance for isoniazid (**C**) and rifampicin (**D**).

After repeated concomitant exposure to rifampicin, susceptibility to isoniazid dropped through lower efficacy (associated with a decrease in E_MAX_, the maximum antimycobacterial effect, of 56.8%) (Table 1 and Figure 1A). Isoniazid resistance emerged later (associated with a decrease in model parameter λ of 32.0%) in the presence of rifampicin (Table 1 and Figure 1C), i.e. the tail in bi-phasic mycobacterial killing curve emerged later. Presence of isoniazid improved rifampicin susceptibility (associated with an increase in E_MAX_ of 42.1%) but rifampicin resistance emerged earlier (associated with an increase in model parameter λ of 58.2%) in the presence of isoniazid (Table 1 and Figure 1B & 1D).

A bi-phasic killing profile was observed, that was more pronounced during the second evolution cycle, and this could consist of different genotypic resistant and phenotypic persistent sub-populations.

### Emergence of genotypic resistance

Whole Genome Sequencing (WGS) of samples from the static kill curve experiments at baseline and at the end of the first and second evolution cycle was used to disentangle emergence of genotypic and phenotypic resistance. Emergence of phenotypic isoniazid resistance (Table 1 and Figure 1C) coincides with appearance of SNPs in genes associated with isoniazid resistance and these accumulate after repeated exposure (Figure 2A). However, isoniazid resistance-associated nucleotide changes occurred stochastically in one out of 12 experiments and with accumulation after repeated exposure in only three out of 12 experiments (Figure 2C). The frequency of isoniazid resistance-associated nucleotide changes was higher in experiments with lower isoniazid exposure and presence of rifampicin did not result in suppression of isoniazid resistance-associated nucleotide changes from emerging. This indicates that isoniazid induced bi-phasic mycobacterial killing can be partly explained by concentration dependent emergence of isoniazid resistant SNPs while the remainder can be attributed to emergence of phenotypic resistance.

**Fig. 2.**
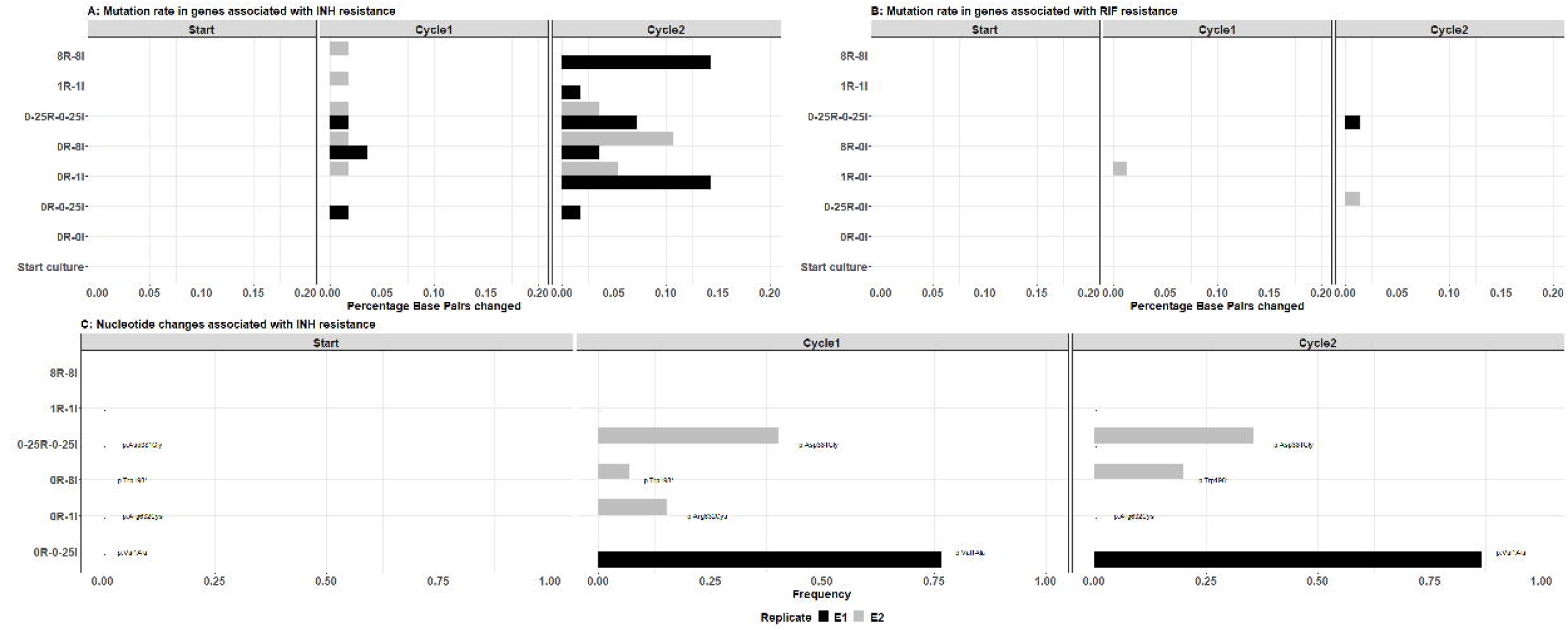
Emergence of isoniazid and rifampicin resistance. Mutation rate for genes associated with isoniazid (**A**) and rifampicin (**B**) resistance. Nucleotide changes associated with isoniazid resistance (**C**).

No rifampicin resistance-associated nucleotide changes were observed and the mutation rate in genes associated with rifampicin resistance was lower when compared to mutation rates in genes associated with isoniazid resistance (Figure 2A & 2B). Consequently, rifampicin induced bi-phasic mycobacterial killing could only be attributed to emergence of phenotypic resistance.

### Impact of dosing regimen

The impact of various dosing strategies was evaluated using the Hollow-Fibre Infection Model (HFIM) and increased dosing regimens, such as Omnie die (OD) bolus dosing at three times the standard dosage or Ter die sumendum (TDS) dosing with standard dosage displayed a tendency of increased bacillary clearance, i.e. linear model slope: 1.61 Mycobacterium Growth Indicator-Tube Time To Positivity (MGIT-TTP)/day or 1.40 MGIT-TTP/day, when compared to O.D. dosing at standard dosage, linear model slope: 1.05 MGIT-TTP/day. Spreading out the dosage by TDS dosing of 1/3 of standard dosage also resulted in increased bacillary clearance, i.e. linear model slope: 1.31 MGIT-TTP/day. However, there was no trend of bi-phasic bacillary clearance for any of the four tested dosing regimens using a HFIM (Figure 3A) and drug-resistant associated SNPs only occurred at the start in one out of four experiments (TDS at 1/3 standard dose) and disappeared over the course of the experiment (Figure 3B). This indicates that emergence of genotypic resistance did not occur at the mimicked dosing regimens.

**Fig. 3:**
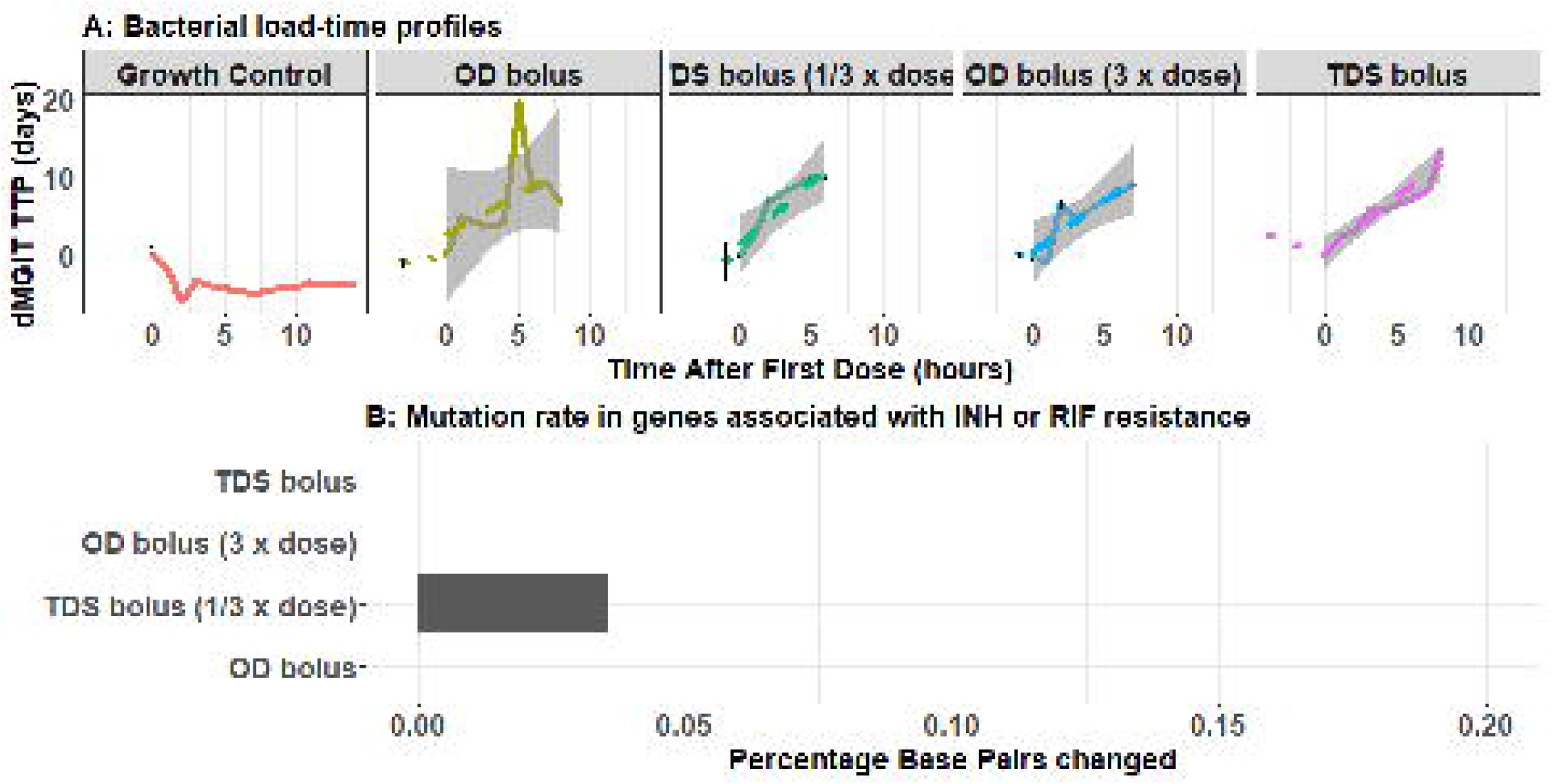
Bacterial killing in the HFIM. Bacterial growth and killing after four different dosing regimens simulated in the HFIM (**A**). Mutation rate for genes associated with isoniazid or rifampicin resistance (**B**).

## DISCUSSION

While investigating pharmacodynamic drug-drug interactions and emergence of resistance, with static kill curve experiments describing two evolution cycles, it was shown that rifampicin resistance was solely phenotypic (i.e. non-specific resistance mechanisms) but isoniazid could be attributed to phenotypic (non-specific) as well as specific genotypic mechnaisms (Figure 1-2). Genotypic isoniazid resistance tends to occur at lower exposures (Figure 2). However, for each of the four tested dosing regimens mimicked in the HFIM, that ranged from standard regimens (*2*) to increased dosing regimens similar to those currently being investigated in clinical trials (*4, 5*), there were no sign of emergence of resistance (Figure 3). Since our findings also displayed antimicrobial exposure dependent bacillary clearance, as observed in patients (*4, 16*), we confirm that pharmacokinetic endpoints such as AUC/MIC and Cmax/MIC, i.e. the main pharmacokinetic endpoints in anti-tuberculosis therapy (*17, 18*), have been attained in all four dosing regimens and that exposures are well above target levels or emergence of resistance.

However, previous studies with isoniazid and rifampicin in the HFIM did report the emergence of at least phenotypic resistance in the first 7 days (*19-21*). A possible explanation for discrepancies in emergence of phenotypic resistance might be quantification methods for bacillary load given that inoculums were similar around 10^6^. Previous studies used Colony Forming Units (CFU’s) as the quantification method, while here we used MGIT-TTP, which is known to have increased sensitivity associated with optimized liquid growth media and detection of oxygen consumption (*22*). Nevertheless, our results show that emergence of genotypic resistance at clinical and investigational dose levels is unlikely provided patients adhere to their treatment regimen. However, more detailed investigations could provide an improved mechanistic understanding of the benefits to be expected from optimization of dosing regimens (*4, 5*), and accounting for genetic polymorphism in genes coding for metabolizing enzymes (*23*). This research would require focus on gene expression, metabolomics and lipidomics in order to dissect and quantify sub-populations by shifts in lipid metabolism, cell wall thickening as well as drug specific responses such as mycolic acid pathway caused by isoniazid and upregulation of drug targets for rifampicin (*7*).

Adherence scenarios were not studied using the HFIM but on the basis of results from the static kill curve experiments it could be concluded that continued exposure to sub- or MIC equivalent drug levels increase the chance of isoniazid resistance emergence. This emphasises the importance of adherence to the treatment (*24*) in order to prevent low isoniazid exposure and consequent emergence of resistance. In this study only two evolution cycles were performed; a larger number of evolution cycles might have also resulted in emergence of rifampicin resistance as the mutation rate to rifampicin resistance is lower (*11-15*) compared to isoniazid (*10, 11*).

In this study we adopted the HFIM to model the dynamic exposure of bacteria to antimicrobials. There are discrepancies between static kill curve and HFIM models and these are likely a result of the experimental design, for example the persistence of the antibiotic in the media. It was confirmed that isoniazid and rifampicin crossed the fibers and not stick to plastics in the HFIM (Fig. S1), differences are therefore probably a result of static experiments being susceptible to chemical degradation of antibiotics and depletion of nutrients. Isoniazid is chemically unstable at 37L and concentrations decrease by over 50% over seven days (*25*). While in the HFIM a new bolus was supplied every 24 or 8 hours, isoniazid was not topped up over the course of a 7-day static kill curve evolution cycle. Emergence of phenotypic and genotypic isoniazid resistance in the static kill curve experiments might therefore also have been exacerbated by low isoniazid levels towards the end of an evolution cycle. Rifampicin has been shown to decrease by over 90% over 7 days at 37□ and likewise phenotypic resistance emerged in static kill curve experiments might also have been facilitated in part by declining rifampicin levels over the course of the experiment. Furthermore, the volumes of the experimental models might have been a confounding factor, the volume of a tube in static kill curve experiments was only 8 mL and not refreshed over the course of a 7-day evolution cycle. While the system volume of the HFIM was 108 mL and refreshed on a continuous basis. The phenotypic resistance that emerged in rifampicin static kill curve experiments might have been stress and not drug associated which could explain the absence of rifampicin genotypic resistance.

Here, we demonstrate that emergence of antimicrobial specific (genotypic) resistance only occurs when antibiotic levels fall below MIC levels, however, MICs are maintained following clinical dosing provided that adherence to the regimen is good. Further work to understand the mechanisms underlying concentration dependent emergence of phenotypic resistance is desirable in order to improve the drug development pipeline and design of novel regimens.

## MATERIALS AND METHODS

### Study design

Static kill curve experiments, to elucidate emergence of resistance defined by phenotype and genotype, comprised three stages; during the first evolution cycle cultures were exposed to various isoniazid and/or rifampicin concentrations for one week, subsequently cultures were regrown in drug free media for three weeks and during the second evolution cycle cultures were exposed to the same drug conditions as the first evolution cycle for another week. This allowed for selection of bacterial sub-population that were able to survive isoniazid and/or rifampicin exposure.

Hollow-fibre infection model experiments, to investigate the impact of dose intervals on resistance emergence, mimicked O.D. oral intake of 600 mg and 300 mg rifampicin and isoniazid, O.D. oral intake of 1,800 mg and 900 mg rifampicin and isoniazid, T.D.S. oral intake of 600 mg and 300 mg rifampicin and isoniazid and T.D.S. oral intake of 200 mg and 100 mg rifampicin and isoniazid.

All static kill curve and hollow-fibre infection model experiments were conducted in biosafety level 3 laboratories.

### Static kill-curve experiments

All drug experiments and growth controls were performed in duplicate at 37°C and ambient air. Isoniazid (40 mg/mL) and rifampicin (83 mg/L) stock solutions were prepared with sterile distilled water directly from the drug vials (Becton Dickinson - MGIT 960 SIRE KIT). Stock solutions were then diluted to drug experiment conditions containing 250, 1,000 or 8,000 ng/mL of isoniazid and 12.5, 50, and 400 ng/mL rifampicin alone or in combination with 10% OADC-supplemented Middlebrook broth 7H9 (Becton Dickinson - BBL MGIT 7ML).

*M. tuberculosis* H37Rv (NCTC 7416, ATCC 9360, obtained from Public Health England culture collections), was incubated in a MGIT Tube containing 7 mL Middlebrook broth 7H9 (Becton Dickinson - BBL MGIT 7ML), supplemented with 10% OADC (Becton Dickinson - MGIT OADC ENRICHMENT 6 VIALS) prior to the first evolution cycle. Evolution cycle one was started once the bacterial load of the master culture reached 10^5.5^ CFU/mL. Each drug experiment or growth control experiment started with inoculating MGIT tubes containing 7 mL Middlebrook broth 7H9 with pellets from the master tube resuspended in 0.5 mL 7H9, 0.8 mL OADC and 0.2 mL drug solution, rending a total volume of 8.5 mL with a bacterial load of ∼ 10^6^ CFU/mL.

The bacteria were separated from media by centrifugation at 2,683 Relative centrifugal force (RCF) for 10 minutes at the end of the first evolution cycle and the regrowth phase of the experiment. Pellets were washed in Sterile phosphate-buffered saline (PBS) and pelleted at 2,683 RCF for 10 minutes twice. Thereafter the pellets were resuspended in the new condition at the start of the second or third stage in volumes as described for initial inocula.

Daily samples of 50 μL were taken for bacterial load quantification and three samples (baseline, after the first and after the second evolution cycle) were taken for Whole Genome Sequencing.

### Hollow-fibre infection model experiments

A cellulosic (C3008, FibreCellSystems Ltd) cartridge was inoculated with 20 mL 10^5.5^ *M. tuberculosis*, strain H37Rv. Incubation followed the identical procedure as per kill-curve experiments. A drug-free incubation phase of two-three days preceded the start of the drug experiments.

Isoniazid and rifampicin concentration-time profiles mimicked unbound concentrations in the granuloma under the assumption that there is equilibrium between unbound drug concentrations in plasma and the granuloma (*26*). Model predictions were obtained with a previously published model, adjusted for 42% and 83% plasma protein binding for isoniazid and rifampicin, respectively (*27*). The following pharmacokinetic properties were mimicked for the OD and TDS standard dosage experiments: isoniazid C_max_ = 3.32 mg/l and rifampicin C_max_ =0.788 mg/l with t_1/2_=4.7h). For the TDS experiment with 1/3 of standard dosing isoniazid C_max_ was 1.06 mg/l and rifampicin C_max_ =0.263 mg/l with a t_1/2_ at 4.7h. The OD experiment at 3 times standard dosing simulated isoniazid C_max_ at 9.50 mg/l and rifampicin C_max_ at 2.36 mg/l with a t_1/2_ at 4.7h. A web application (https://pkpdia.shinyapps.io/hfs_app/) (*26*) was subsequently used to convert secondary pharmacokinetic parameter estimates, CMAX/C0 and t1/2, into pump settings at a system volume of 108 mL (central reservoir, 50 mL; intracapillary space and tubing, 44 mL; and extracapillary space, 14 mL). First-order absorption of the drugs, to mimic oral absorption characteristics, was omitted and replaced by bolus administration in the central reservoir in these experiments to avoid complex experimental setups.

Growth control and drug experiments lasted for 14 and 7 days, respectively. Drug concentrations in the hollow-fibre medium were not measured during the experiments but pump-settings were validated using bacteria free experiments to ensure *in-vivo* mimicking pharmacokinetic profiles were simulated *in-vitro* (Fig. S1).

Daily samples of 50 μL were taken for bacterial load quantification and two samples, at baseline at the end of the drug experiments, were taken for Whole Genome Sequencing.

### Bacterial load quantification

Fifty μL samples from kill-curve and hollow-fibre infection model experiments were inoculated in fresh MGIT tubes containing 7 mL Middlebrook broth 7H9 with 0.8 mL OADC. MGIT-TTP was used as measure of bacterial load (*22*).

### Whole Genome Sequencing

The CTAB method was used to extract genomic DNA and Qubit dsDNA kits (Life Technologies) were used to quantify DNA prior to sequencing (*28*). WGS for start culture and evolution cycle two samples were performed using an Illumina HiSeq platform and samples from the first evolution cycle and all Hollow-fibre infection model experiments were performed using the Illumina NextSeq platform. WGS was performed following the manufacturers’ instructions and local validated protocols and results have been deposited (Table S1).

### Drug quantification

Pharmacokinetic validation experiments were performed under identical conditions as described above, apart from that the cartridge was not inoculated with *M. tuberculosis* and broth was not supplemented with OADC. Broth samples (2 mL) were taken from the central reservoir 0, 3, 6, 8 and 23 hours post bolus dose, and from the extra capillary space 3 and 24 hours post bolus dose. Samples were kept in -80°C until analysis using ultra-high-performance liquid chromatographic-tandem mass spectrometric detection. Details on the analytical method have been published previously (*26*).

### Pharmacodynamic modelling

A compartmental model was developed to describe MGIT-TTP data from time kill curve experiments (Fig. S2 & Data file S1). Model estimates were computed using the SAEM estimation method in nlmixr 2.0.4 on a Windows 10 operating system. Minus twice the log likelihood of the data was used as objective function value (OFV) and a drop in OFV of at least 3.84 (P=0.05) was considered to improve the model’s ability to fit the data with statistical significance after inclusion of one degree of freedom to a nested hierarchical model. Assessment of model performance was further supported by goodness-of-fit diagnostics including observation-population predictions, observations-individual predictions, Normalised Prediction Distribution Errors (NPDE)-population predictions and NPDE-time.

Baseline MGIT-TTP at experiment level (*P*_*i*_) was estimated using a typical baseline MGIT-TTP (*θ*_*TV*_) and a deviation from the typical baseline MGIT-TTP described by between experiment variability (η) (Eq. 1).

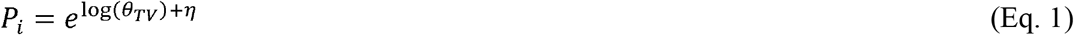

Data from the growth control experiments were used to develop a log-growth model (Eq. 2) using the parameters net growth 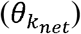 and maximum carrying capacity 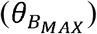. The parameters 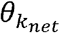 and 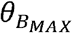 followed log-normal distribution for between experiment variability as described in Eq. 1. Data from re-growth experiments was subsequently included and predictive performance of the models was evaluated.

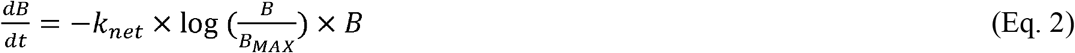

Isoniazid and rifampicin anti-mycobacterial effects were parameterised (Eq. 3) using maximum drug efficacy 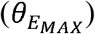 and potency 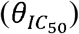, i.e. the concentration at which half-maximum inhibition is achieved, with C being the isoniazid or rifampicin concentration and (*θ*_*γ*_) being the shape factor.

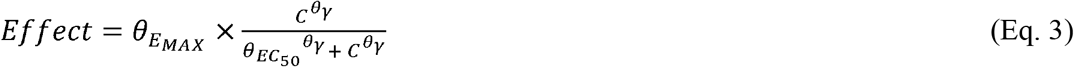

Loss of susceptibility to isoniazid and or rifampicin over the course of the experiment was parameterised using *θ*_*BETA*_, i.e. magnitude of susceptibility loss, and *θ*_*λ*_, i.e. onset of susceptibility loss as a function of time (Eq. 4).

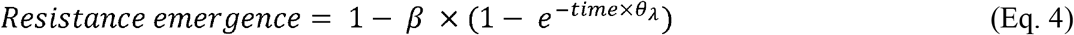

Drug combinations were evaluated as additive drug effects at first, i.e. 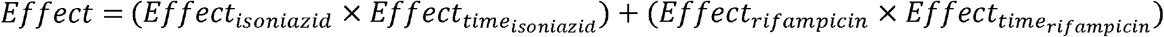, and subsequently the Bliss independence model was tested and retained (*29*), i.e. 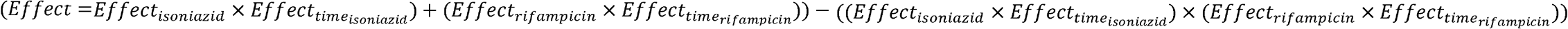. Synergism and antagonism were evaluated as categorical effects for presence of drugs in combination and a drop in OFV of at least 3.84 (P=0.05) was considered to improve the model’s ability to fit the data with statistical significance in stepwise covariate model building (*30*).

### Whole Genome Sequence analysis

Analysis of WGS data was divided in two steps. First, all mutations relative to the H37Rv reference genome were determined for 33 genes of interest associated with resistance to rifampicin, isoniazid, ethambutol, pyrazinamide, streptomycin, ethionamide, amikacin, capreomycin, kanamycin, fluoroquinolones, para-aminosalicylic acid, cycloserine, linezolid, bedaquiline, clofazimine and delamanid (*rpoB, rpoC, fabG1, inhA, katG, kasA, ahpC, embR, embC, embA, embB, rpsA, pncA, panD, rpsL, gid, rrs, ethA, ethR, tlyA, eis, gyrB, gyrA, folC, ribD, thyX, thyA, ald, alr, rplC, rrl, Rv0678* and *fbiA*). Bcftools (v. 1.11), bwa (v. 0.7.12) and samtools (v. 1.9) were used for variant calling using the number of forward reference, reverse reference, forward non-reference, reverse non-reference alleles. The percentage of base pairs changed within genes, were stratified by genes associated with isoniazid and/or rifampicin resistance. Only base pairs with ≥15% changed to the H37Rv reference genome were used to calculate the percentage of base pairs changed within genes.

Tb-profiler (v. 3.0.1) was used to determine the SNPs by nucleotide changes (*31*). Results were stratified by drug resistant nucleotide changes, i.e. isoniazid and rifampicin associated resistance, and other resistance that associate with any other of the aforementioned drugs.

## Supporting information

PKPD model

## Supplementary Materials

Fig. S1. Simulated (grey dashed line) versus observed pharmacokinetic profile in central reservoir (circles) and extra capillary space (triangles).

Fig. S2. Basic Goodness of Fitness plots Pharmacodynamic model. Grey and blue lines represent the line of identity and local polynomial regression fitting, respectively.

Table S1. Summary of WGS data.

Data file S1. Pharmacodynamic model, code and data.

## Acknowledgments

None

## Funding

UKRI Medical Research Council Skills Development Fellowship MR/P014534/1 (FK)

## Author contributions

Conceptualization: FK, TDM

Methodology: FK, AW, NS, TDM

Investigation: FK, JO, LP, KK, ZS

Visualization: FK

Funding acquisition: FK

Project administration: FK

Supervision: FK

Writing – original draft: FK

Writing – review & editing: FK, JO, LP, ZS, KK, AW, NS, TDM

## Competing interests

Authors declare that they have no competing interests.

## Data and materials availability

Accession numbers to WGS data are on an ENA project (Table S1) and the pharmacodynamic drug-drug interaction model is attached as Data file S1.

## Notes

### Competing Interest Statement

The authors have declared no competing interest.

