## Supplement figures and tables for "Understanding emergence of antimycobacterial dose dependent resistance"

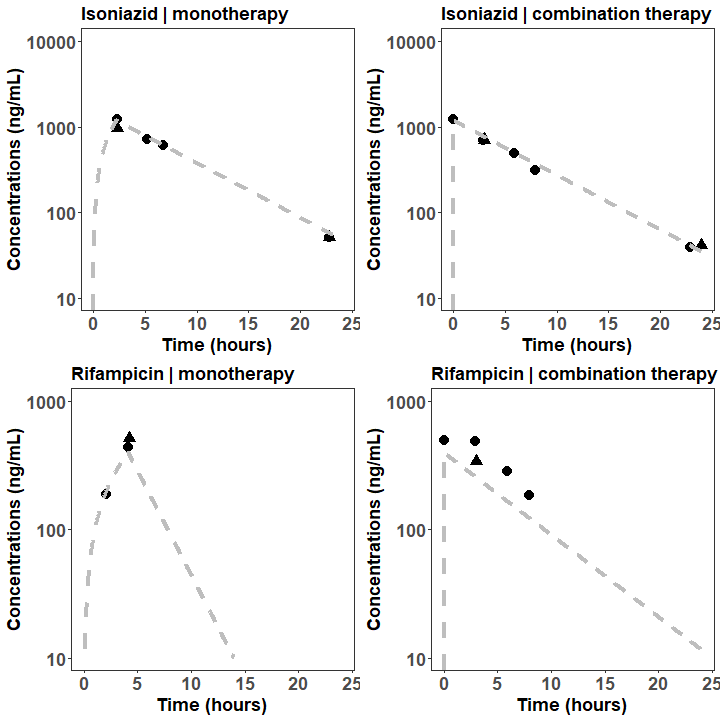

Fig. S1. Simulated (grey dashed line) versus observed pharmacokinetic profile in central reservoir (circles) and extra capillary space (triangles).

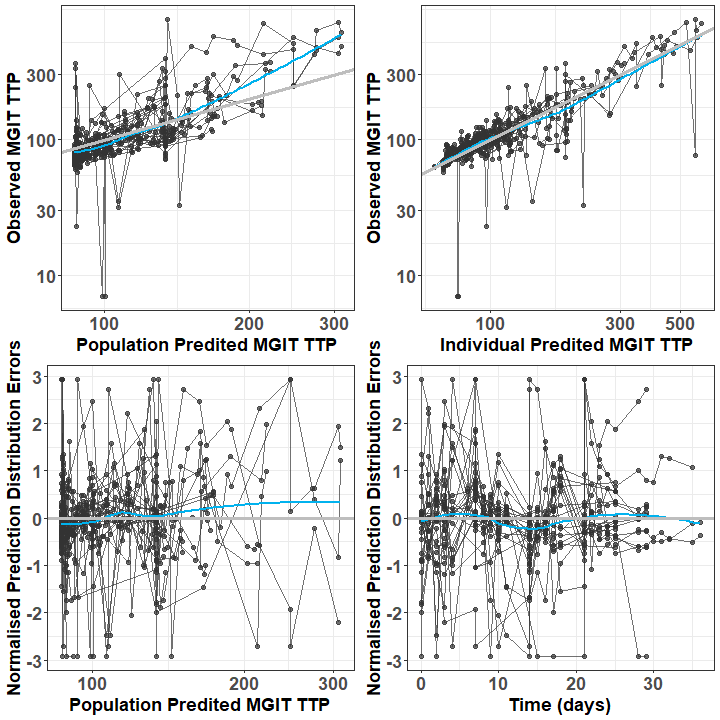

Fig. S2. Basic Goodness of Fitness plots Pharmacodynamic model. Grey and blue lines represent the line of identity and local polynomial regression fitting, respectively.

Table S1. Summary of WGS data.

| **id** | **project_title** | **scientific_name** | **strain** | **sample_alias** | **sample_description** |
| --- | --- | --- | --- | --- | --- |
| ERS12274705 | PRJEB53040 | Mycobacterium tuberculosis | H37Rv | E1_0-25R-0-25I-2 | 1/4 MIC RIF and 1/4 MIC INH, end of evolution cycle 2, replicate 1 |
| ERS12274706 | PRJEB53040 | Mycobacterium tuberculosis | H37Rv | E1_0-25R-0I-2 | 1/4 MIC RIF and 0 MIC INH, end of evolution cycle 2, replicate 1 |
| ERS12274707 | PRJEB53040 | Mycobacterium tuberculosis | H37Rv | E1_0R-0-25I-2 | 0 MIC RIF and 1/4 MIC INH, end of evolution cycle 2, replicate 1 |
| ERS12274708 | PRJEB53040 | Mycobacterium tuberculosis | H37Rv | E1_0R-0I-2 | 0 MIC RIF and 0 MIC INH, end of evolution cycle 2, replicate 1 |
| ERS12274709 | PRJEB53040 | Mycobacterium tuberculosis | H37Rv | E1_0R-1I-2 | 0 MIC RIF and 1 MIC INH, end of evolution cycle 2, replicate 1 |
| ERS12274710 | PRJEB53040 | Mycobacterium tuberculosis | H37Rv | E1_0R-8I-2 | 0 MIC RIF and 8 MIC INH, end of evolution cycle 2, replicate 1 |
| ERS12274711 | PRJEB53040 | Mycobacterium tuberculosis | H37Rv | E1_1R-0I-2 | 1 MIC RIF and 0 MIC INH, end of evolution cycle 2, replicate 1 |
| ERS12274712 | PRJEB53040 | Mycobacterium tuberculosis | H37Rv | E1_1R-1I-2 | 1 MIC RIF and 1 MIC INH, end of evolution cycle 2, replicate 1 |
| ERS12274713 | PRJEB53040 | Mycobacterium tuberculosis | H37Rv | E1_8R-0I-2 | 8 MIC RIF and 0 MIC INH, end of evolution cycle 2, replicate 1 |
| ERS12274714 | PRJEB53040 | Mycobacterium tuberculosis | H37Rv | E1_8R-8I-2 | 8 MIC RIF and 8 MIC INH, end of evolution cycle 2, replicate 1 |
| ERS12274715 | PRJEB53040 | Mycobacterium tuberculosis | H37Rv | E1_Start-culture-2 | Start culture, replicate 1 |
| ERS12274716 | PRJEB53040 | Mycobacterium tuberculosis | H37Rv | E2_0-25R-0-25I-2 | 1/4 MIC RIF and 1/4 MIC INH, end of evolution cycle 2, replicate 2 |
| ERS12274717 | PRJEB53040 | Mycobacterium tuberculosis | H37Rv | E2_0-25R-0I-2 | 1/4 MIC RIF and 0 MIC INH, end of evolution cycle 2, replicate 2 |
| ERS12274718 | PRJEB53040 | Mycobacterium tuberculosis | H37Rv | E2_0R-0-25I-2 | 0 MIC RIF and 1/4 MIC INH, end of evolution cycle 2, replicate 2 |
| ERS12274719 | PRJEB53040 | Mycobacterium tuberculosis | H37Rv | E2_0R-0I-2 | 0 MIC RIF and 0 MIC INH, end of evolution cycle 2, replicate 2 |
| ERS12274720 | PRJEB53040 | Mycobacterium tuberculosis | H37Rv | E2_0R-1I-2 | 0 MIC RIF and 1 MIC INH, end of evolution cycle 2, replicate 2 |
| ERS12274721 | PRJEB53040 | Mycobacterium tuberculosis | H37Rv | E2_0R-8I-2 | 0 MIC RIF and 8 MIC INH, end of evolution cycle 2, replicate 2 |
| ERS12274722 | PRJEB53040 | Mycobacterium tuberculosis | H37Rv | E2_1R-0I-2 | 1 MIC RIF and 0 MIC INH, end of evolution cycle 2, replicate 2 |
| ERS12274723 | PRJEB53040 | Mycobacterium tuberculosis | H37Rv | E2_1R-1I-2 | 1 MIC RIF and 1 MIC INH, end of evolution cycle 2, replicate 2 |
| ERS12274724 | PRJEB53040 | Mycobacterium tuberculosis | H37Rv | E2_8R-0I-2 | 8 MIC RIF and 0 MIC INH, end of evolution cycle 2, replicate 2 |
| ERS12274725 | PRJEB53040 | Mycobacterium tuberculosis | H37Rv | E2_8R-8I-2 | 8 MIC RIF and 8 MIC INH, end of evolution cycle 2, replicate 2 |
| ERS12274726 | PRJEB53040 | Mycobacterium tuberculosis | H37Rv | E2_Start-culture-2 | Start culture, replicate 2 |
| ERS12274727 | PRJEB53040 | Mycobacterium tuberculosis | H37Rv | 0R-0I-7 | 0 MIC RIF and 0 MIC INH, end of evolution cycle 1, replicate 2 |
| ERS12274728 | PRJEB53040 | Mycobacterium tuberculosis | H37Rv | 0R-1-4I-1 | 0 MIC RIF and 1/4 MIC INH, end of evolution cycle 1, replicate 2 |
| ERS12274729 | PRJEB53040 | Mycobacterium tuberculosis | H37Rv | 0R-1-4I-9 | 0 MIC RIF and 1/4 MIC INH, end of evolution cycle 1, replicate 1 |
| ERS12274730 | PRJEB53040 | Mycobacterium tuberculosis | H37Rv | 0R-1I-5 | 0 MIC RIF and 1 MIC INH, end of evolution cycle 1, replicate 2 |
| ERS12274731 | PRJEB53040 | Mycobacterium tuberculosis | H37Rv | 0R-8I-8 | 0 MIC RIF and 8 MIC INH, end of evolution cycle 1, replicate 1 |
| ERS12274732 | PRJEB53040 | Mycobacterium tuberculosis | H37Rv | 0R-8I-11 | 0 MIC RIF and 8 MIC INH, end of evolution cycle 1, replicate 2 |
| ERS12274733 | PRJEB53040 | Mycobacterium tuberculosis | H37Rv | 1-4R-1-4I-2 | 1/4 MIC RIF and 1/4 MIC INH, end of evolution cycle 1, replicate 2 |
| ERS12274734 | PRJEB53040 | Mycobacterium tuberculosis | H37Rv | 1-4R-1-4I-10 | 1/4 MIC RIF and 1/4 MIC INH, end of evolution cycle 1, replicate 1 |
| ERS12274735 | PRJEB53040 | Mycobacterium tuberculosis | H37Rv | 1R-1I-6 | 1 MIC RIF and 1 MIC INH, end of evolution cycle 1, replicate 2 |
| ERS12274736 | PRJEB53040 | Mycobacterium tuberculosis | H37Rv | 8R-0I-3 | 8 MIC RIF and 0 MIC INH, end of evolution cycle 1, replicate 2 |
| ERS12274737 | PRJEB53040 | Mycobacterium tuberculosis | H37Rv | 8R-8I-4 | 8 MIC RIF and 8 MIC INH, end of evolution cycle 1, replicate 2 |
| ERS12274738 | PRJEB53040 | Mycobacterium tuberculosis | H37Rv | 0R-0I-15 | 0 MIC RIF and 0 MIC INH, end of evolution cycle 1, replicate 1 |
| ERS12274739 | PRJEB53040 | Mycobacterium tuberculosis | H37Rv | 0R-1I-13 | 0 MIC RIF and 1 MIC INH, end of evolution cycle 1, replicate 1 |
| ERS12274740 | PRJEB53040 | Mycobacterium tuberculosis | H37Rv | 1-4R-0I-14 | 1/4 MIC RIF and 0 MIC INH, end of evolution cycle 1, replicate 1 |
| ERS12274741 | PRJEB53040 | Mycobacterium tuberculosis | H37Rv | 1R-0I-16 | 1 MIC RIF and 0 MIC INH, end of evolution cycle 1, replicate 2 |
| ERS12274742 | PRJEB53040 | Mycobacterium tuberculosis | H37Rv | 8R-0I-12 | 8 MIC RIF and 0 MIC INH, end of evolution cycle 1, replicate 1 |
| ERS12274743 | PRJEB53040 | Mycobacterium tuberculosis | H37Rv | Day0-Exp1 | Start 1/3 x standard dose TDS HFIM experiment |
| ERS12274744 | PRJEB53040 | Mycobacterium tuberculosis | H37Rv | Day0-Exp2 | Start 3 x standard dose OD HFIM experiment |
| ERS12274745 | PRJEB53040 | Mycobacterium tuberculosis | H37Rv | Day6-Exp1 | End 1/3 x standard dose TDS HFIM experiment |
| ERS12274746 | PRJEB53040 | Mycobacterium tuberculosis | H37Rv | Day7-Exp2 | End 3 x standard dose OD HFIM experiment |
| ERS12274747 | PRJEB53040 | Mycobacterium tuberculosis | H37Rv | H37Rv-1 | 1/4 MIC RIF and 0 MIC INH, end of evolution cycle 1, replicate 2 |
| ERS12274748 | PRJEB53040 | Mycobacterium tuberculosis | H37Rv | H37Rv-2 | 1 MIC RIF and 0 MIC INH, end of evolution cycle 1, replicate 1 |
| ERS12274749 | PRJEB53040 | Mycobacterium tuberculosis | H37Rv | H37Rv-3 | 8 MIC RIF and 8 MIC INH, end of evolution cycle 1, replicate 1 |
| ERS12274750 | PRJEB53040 | Mycobacterium tuberculosis | H37Rv | H37Rv-4 | 1 MIC RIF and 1 MIC INH, end of evolution cycle 1, replicate 1 |
| ERS12274751 | PRJEB53040 | Mycobacterium tuberculosis | H37Rv | Frank_HFA_OD_23AUG2019_end-355082158 | End standard dose OD HFIM experiment |
| ERS12274752 | PRJEB53040 | Mycobacterium tuberculosis | H37Rv | Frank_HFB_TDS_28NOV2019_start-355004492 | Start standard dose TDS HFIM experiment |
| ERS12274753 | PRJEB53040 | Mycobacterium tuberculosis | H37Rv | Frank_HFC_TDS_10DEC2019_end-355119001 | End standard dose TDS HFIM experiment |
